# Effect of larval diet on adult immune function is contingent upon selection history and host sex in *Drosophila melanogaster*

**DOI:** 10.1101/2022.03.03.482770

**Authors:** Aparajita Singh, Aabeer Kumar Basu, Nitin Bansal, Biswajit Shit, Tejashwini Hegde, Nagaraj Guru Prasad

**Affiliations:** Department of Biological Sciences, Indian Institute of Science Education and Research Mohali, India; School of Biological Sciences, University of Nebraska-Lincoln, NE 68588, USA; Department of Biology, Ashoka University, Sonipat, India; Institut für Populationsgenetik, Vetmeduni Vienna, Veterinärplatz 1, 1210, Vienna, Austria

**Keywords:** *Drosophila melanogaster*, experimental evolution, bacterial pathogens, poor nutrition, life-history trade-offs

## Abstract

Mounting an immune response requires a considerable energy investment by the host. This makes expression of immune phenotypes susceptible to changes in availability of resources. There is ample evidence in scientific literature to suggest that hosts become more vulnerable to infection by pathogens and parasites when access to nutrition is limited. Using populations of *Drosophila melanogaster* experimentally evolved to better survive bacterial infections, we explore if host selection history influences host response to resource deprivation in terms of immune function. We find that when reared on a suboptimal diet (both in larval and adult stages), adult flies from evolved populations are still more immune to infections compared to flies from control populations. Furthermore, we observe a sex-dependent effect of interaction between selection history and diet on immune function. We thus conclude that immune function of hosts adapted to pathogen challenge is less affected by resource limitation compared to non-adapted hosts.

## 1. INTRODUCTION

Expression of immune phenotypes is governed by trade-offs, both evolutionary and physiological (Sheldon and Verhulst 1996). Evolutionary trade-offs stem from antagonistic pleiotropy or linkage disequilibrium, with more immune-competent genotypes having sub-optimal fitness in terms of other organismal traits such as reproduction (Schmid-Hempel 2003). Physiological trade-offs are driven by differential resource allocation between immune function and other organismal traits; increased investment towards immune defense compromises other life history traits of the individual organism, and vice versa (Lochmiller and Deerenberg 2000). The resource allocation towards immune function can be plastic (depending on environmental factors like exposure to pathogens, availability of resources, etc.) or developmentally pre-determined. Additionally, the cost of investment towards immune function in terms of its negative effects on other traits often manifest only when the individual organism is subjected to infection (McKean et al. 2008, Lazzaro and Little 2009).

Studying correlated responses to selection in controlled evolution set-ups is an easy method to elucidate trade-offs between different organismal traits. Such set-ups have repeatedly been used to study evolution of defense against parasites and pathogens, and correlated evolution of other life-history traits. *Drosophila melanogaster* populations selected for increased defense against larval parasitoid show reduced capacity of intra-specific competition (Kraaijeveld and Godfray 1997, Fellowes et al. 1998). Similarly, *Drosophila* flies selected for better resistance against bacterial pathogen *Pseudomonas aeruginosa* have reduced egg viability and adult life-span (Ye et al. 2009). Populations of Indian meal moth, *Plodia interpunctella*, selected for increased resistance to granulosis virus exhibit increased development time and reduced egg viability (Boots and Begon 1993). Red flour beetles, *Tribolium casteneum*, populations selected for increased immune defense against bacterium *Bacillus thuringiensis* exhibit reduced egg and juvenile viability (Prakash et al. 2022).

Contrary to expectations, associated life-history trade-offs are not always observed in experimental evolution studies. For example, both Faria et al. (2015) and Gupta et al. (2016) selected adult *Drosophila melanogaster* flies for resistance against the same pathogenic bacteria, *Pseudomonas entomophila*, and did not find observable trade-offs with any of the measured life-history traits. Additionally, in *Drosophila melanogaster* populations where larval competitive ability trade-offs with parasitoid defense, no trade-off is observed with respect to fecundity, egg viability, and starvation resistance (Fellowes et al. 1998).

Laboratory populations live in an environment with ample access to resources, and this might be the reason why trade-offs are not always observed in laboratory experimental evolution studies (Harshman and Hoffman 2000). It has been often argued that trade-offs only manifest under stressful conditions (Reznick 1985, Stearns 1989, Marden et al. 2003). In fact, the trade-off between parasitoid defense and intra-specific competitive ability in *Drosophila melanogaster* is only observed when resources are scarce (Kraaijeveld and Godfray 1997, Fellowes et al. 1998). Excess resources are also known to help ameliorate reproduction immunity trade-off in *Drosophila melanogaster* (McKean and Nunney 2005) and *Tenebrio molitor* (Ponton et al. 2010). Therefore, one way to identify immune function associated trade-offs may be to assess immunity and life-history traits under resource limited conditions.

There is ample evidence that host organisms exposed to poor nutrition suffer from reduced immune defense and increased susceptibility to pathogens. Starvation reduces phenoloxidase activity in mealworm beetle, *Tenebrio molitor* (Siva-Jothy and Thompson 2002). Probability of survival till adulthood for mosquito (*Aedes aegypti*) larvae infected with microsporidian parasite (*Vavria culicis*) increases with increase in food availability (Bedhomme et al. 2004). Tobacco hornworm, *Manduca sexta*, when raised on non-native host plants have reduced melanization and encapsulation capacity (Diamond and Kingsolver 2011). Limiting access to nutrition can alter the functionality of different components of the host immune system, instead of universal downregulation, making the effect of malnutrition on host immunity mechanism and pathogen specific (Adamo et al. 2016). Reducing yeast content in adult diet increases the susceptibility of *Drosophila melanogaster* females to infection with *Pseudomonas entomophila* (Kutzer et al. 2018), but not to infection with *Escherichia coli* or *Lactococcus lactis* (Kutzer and Armitage 2016). In addition to total nutrition availability, host immune function is also affected by changes in specific components of the diet (Cotter et al. 2011). For example, low protein/high carbohydrate diets enhance survival of *Drosophila melanogaster* when infected with *Micrococcus luteus* (Ponton et al. 2020) and of fruit fly *Bactrocera tryoni* when infected with *Serratia marcescens* (Dinh et al. 2019). Similarly, burying beetle *Nicrophorus vespilloides* can survive infection with bacterium *Photorhabdus luminiscens* when fed a high fat/low protein diet (Miller and Cotter 2018). Nutrition can also affect immune function indirectly by affecting the physiological state of the host organism (Diamond and Kingsolver 2011).

Reduction in resources (in the form of reduced access to nutrition), instead of unmasking trade-offs, can also lead to improvement of immune function, in a pathogen specific manner. Ayres and Schneider (2009) raised *Drosophila melanogaster* flies on poor diet as larvae and found the adults to be more resistant to infection with *Salmonella typhimurium*, while being more susceptible to *Listeria monocytogenes*; resistance to *Enterococcus faecalis* remained unchanged. Fly mutants that feed less also exhibit identical patterns of resistance (Ayres and Schneider 2009). Reduced feeding in response to infection is observed in many animal species, but it is unclear if this is an adaptive strategy on the part of the host to defend against infections (Hite et al. 2020). In response to infection, hosts may also modify their choice of food substrate in order to accommodate their immediate dietary requirements (Abbott 2014). For example, bacterial infection in *Drosophila melanogaster* (Ponton et al. 2020) and *Bactrocera tryonni* (Dinh et al. 2019) has also been shown to shift diet choice of flies towards more carbohydrate rich food; general reduction of feeding in *Drosophila* infected with both bacteria and fungi has also been reported (Bashir-Tanoli and Tinsley 2014). Contrary to this, *Sodoptera littoralis* when infected with nucleopolyhedrovirus (Lee et al. 2006) and *Sodoptera exempta* when infected with *Bacillus subtilis* (Povey et al. 2009) prefer high protein diets. An additional source of complexity is that, since pathogens and parasites are dependent on host to acquire resources for their own proliferation, limiting host’s access to nutrition can negatively impact within-host pathogen growth and thereby bias infection outcome in favour of the host (Cressler et al. 2014, Pike et al. 2019).

In this study, we explore how hosts evolved to be more immune to bacterial pathogens respond to scarcity of resources, in terms of immune function and life-history traits. We experimentally evolved two sets of replicate *Drosophila melanogaster* populations to better survive infection with pathogenic bacteria. One set of four replicate populations (with ancestrally paired controls) were selected against a Gram-negative bacterium, *Pseudomonas entomophila* (Gupta et al. 2016), while the another set of four replicate populations (with ancestrally paired controls) were selected against a Gram-positive bacterium, *Enterococcus faecalis* (Singh et al. 2021, preprint). Both selection regimes were derived off the same ancestral baseline population (Figure 1, see METHODS for more details). Using these selection regimes, we tested if rearing on a poor larval diet affected the immune function of adult flies of each selection regime, when infected with their native pathogen. Post-infection survival was used as a proxy of immune function in these experiments. Additionally, we tested if poor larval diet intensifies the trade-off between immune function and life-history traits in the selected populations. Our results indicate that diet and selection history interact to determine post-infection survival of hosts. Additionally, the interactive effect of diet and selection history is not consistent across both sexes.

**Figure 1.**
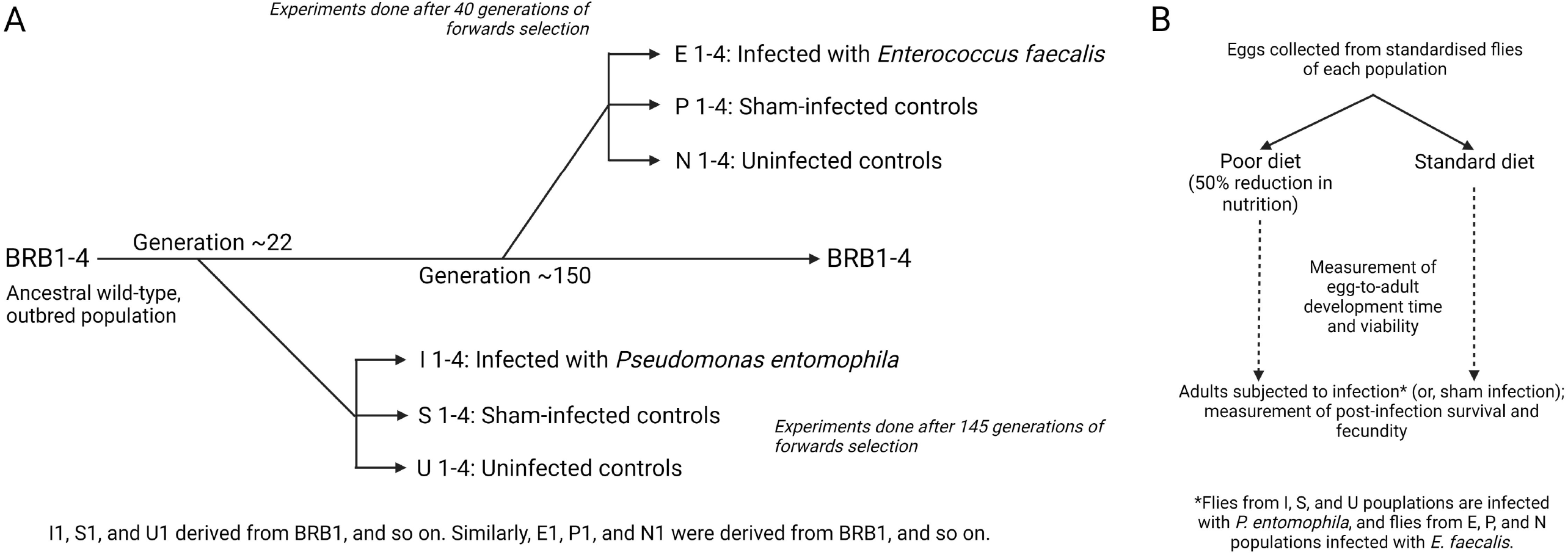
Population history and experimental design. (a) Ancestry of populations used to study the effect of standard vs. poor larval diet on adult immune function in flies, and (b) outline of experimental design.

## 2. MATERIALS AND METHODS

Two selection regimes of *Drosophila melanogaster*, each selected for better survivorship post-infection with a different entomopathogenic bacteria, were used in this study. The Blue Ridge Baseline (BRB) populations were used to derive both selection regimes.

### 2.1 Ancestral populations

Blue Ridge Baseline (BRB) is a set of four independent, large, outbred populations of *Drosophila melanogaster*, referred to as BRB 1-4 (Singh et al 2015). Each replicate population has a census size of 2800 adults and is maintained on a 14-day discrete generation cycle on standard banana-jaggery-yeast food, at 25 °C on a 12:12 light-dark (LD) cycle at 50-60% relative humidity (RH). Larvae are reared at a density of ~70 eggs per glass vial (25 mm diameter × 90 mm height) containing 6-8 mL of standard banana-jaggery-yeast medium (banana 205g, barley flour 25g, jaggery 35g, yeast 36g, agar 12.4g, ethanol 45ml, water 1180ml, and p-hydroxymethyl benzoate 2.4g per litre of standard banana-jaggery-yeast food). There are 40 such vials per population. Almost all flies eclose by the 12^th^ day post egg laying (PEL) and are transferred to plexiglass cages (25 cm length x 20 cm width x 15 cm height) provided with a 90 mm diameter Petri plate containing standard banana-jaggery-yeast food. On the 14th day PEL, a fresh food plate is provided for 18 hours for oviposition. These eggs are used to start the next generation.

### 2.2 Populations selected for better survival post systemic infection with *Enterococcus faecalis* (*E. faecalis*): EPN regime

Selection regime for better survival following systemic infection with entomopathogenic bacteria *Enterococcus faecalis* was started after 150 generations of laboratory adaptation of BRB populations (for details refer Singh et al. 2021, preprint). Briefly, from each replicate population of BRB1-4, three populations were derived: (a) E1-4, infected with *<u>E</u>nterococcus faecalis*, (b) P1-4, <u>p</u>ricking control, and (c) N1-4, <u>n</u>ormal control. Altogether, there were 12 populations in the EPN selection regime: E1-4, P1-4, and N1-4. Populations bearing the same numeral had a more recent common ancestor. For example, E1, P1, and N1 (derived from BRB1) were more closely related to each other than any of them is to E2, P2, and N2 (derived from BRB2) etc. Populations belonging to each block (E1, P1, and N1 constitute block 1, and so on) were handled together on the same day, during both population maintenance and experiments, and were treated as statistical blocks.

At start of each generation, eggs are collected for each population, at a density of 70±10 per vial (25 mm diameter × 90 mm height) containing 6-8 ml of standard banana-jaggery-yeast food; 10 vials for each population. These vials are incubated at standard laboratory conditions as described in Section 2.1. By 9-10^th^ day PEL 95% of the flies eclose. By the 12^th^ day PEL, all of them were mature and had mated at least once. Till this point, all populations are handled in an identical manner.

On the 12^th^ day PEL, E1-4 populations are infected with *E. faecalis*. From each rearing vial, randomly 20 females and 20 males were infected with the help of Minutien pin (0.1 mm, Fine Science Tools, USA) dipped in *E. faecalis* bacterial suspension (see section 2.4) and pricked on the thorax under light CO_2_ anaesthesia. From 10 such vials, total of 200 females and 200 males were infected. After infection, flies were transferred to a plexiglass cage (14 cm length x 16 cm width x 13 cm height) having food plate (60 mm Petri plate in diameter). Fresh food plate was provided every alternate day. Fifty percent of the infected flies would die within 96 hours of infection with *E. faecalis*. Post 96 hours, day 16 PEL, oviposition plates were provided for 18 hours to the population cage to collect eggs for the next generation.

Similarly, on the 12^th^ day PEL, P1-4 populations were pricked with Minutien pin dipped in sterile 10 mM MgSO_4_ buffer under light CO_2_ anaesthesia. From each rearing vial, randomly 10 females and 10 males were pricked. So, total of 100 females and 100 males were sham infected per block. There was negligible mortality (1-2%) post sham infection. Rest of the handling was identical to E1-4.

Handling of N1-4 populations were also similar to E1-4, except, here 10 females and 10 males per vial (total 100 females and 100 males per block) were randomly sorted under light CO_2_ anaesthesia. There was no mortality in N1-4 populations.

Therefore, on day 16, about 100 females and males were present in each population which contributed to the next generation. Thus, EPN selection regime is maintained on a 16-day discrete generation cycle.

### 2.3 Populations selected for better survival post systemic infection with *Pseudomonas entomophila* (*P. entomophila*): IUS regime

Similar to the EPN regime, the IUS regime was derived from BRB1-4 populations after 22 generations of laboratory adaptation as previously described in Gupta et al. (2016). From each replicate population of BRB three populations were derived: (a) I1-4, infected with *Pseudomonas entomophila*, (b) S1-4, <u>s</u>ham-infected control, and (c) U1-4, <u>u</u>ninfected.

The maintenance of IUS regime is identical to that of the EPN regime, except that (a) IUS regime was started from BRB populations after 22 generations of lab adaptation while EPN regime after 150 generations, (b) I flies are infected with Gram-negative bacteria *Pseudomonas entomophila* and E flies with Gram-positive bacteria *Enterococcus faecalis*, (c) for each block of the I1-4 150 females and 150 males whereas for E1-4 200 females and 200 males are infected every generation, (d) peak mortality window, for I is 20 hours to 60 hours and for E is 18 hours to 48 hours, (e) for I1-4 ~33% and for E1-4 50% of the infected flies would die within 96 hours of infection.

### 2.4 Bacterial culture and infection procedure

The bacteria used in the study are *Enterococcus faecalis* (grown at 37 □C) and *Pseudomonas entomophila* (grown at 27 □C). E populations are infected with *E. faecalis*, and I populations with *P. entomophila*. The bacterial stocks are maintained as 17% glycerol stocks frozen at −80 °C. Primary culture of the bacteria is obtained by inoculating a stab of glycerol stock in 10 ml lysogeny broth (Luria-Bertani-Miller, HiMedia) and incubating it overnight at appropriate temperature with continuous mixing at 150 RPM. To establish secondary culture, fresh 10 ml lysogeny broth is inoculated with 100 ul of the overnight culture; incubated as mentioned above till desired turbidity (OD_600_ = 1.0-1.2) is reached. This secondary culture is centrifuged to obtain bacterial pellets which in turn is resuspended in sterile MgSO_4_ buffer (10 mM) to obtain the required optical density (OD_600_). Flies are infected (either during selection protocol or experimental infections) by pricking them on the thorax with a 0.1 mm Minutien pin (Fine Scientific Tools, USA) dipped in the bacterial suspension under light CO_2_ anaesthesia. Sham-infections are carried out similarly, except with a pin dipped in sterile MgSO_4_

### 2.5 Pre-experiment standardization

Prior to any experiment, flies of the selection regimes are reared for a generation under common laboratory conditions. This is done to account for any non-genetic parental effects (Rose 1984), and flies thus generated are called standardized flies. To generate standardized flies, eggs were collected from flies of all the populations at a density of 60-80 eggs per vial; 10 such vials were established per population. The vials were incubated under standard laboratory conditions. On day 12 post egg laying (PEL), by which time almost all the flies would have eclosed, the adults were transferred to plexiglass cages (14×16×13 cm^3^) with food plates (Petri plates, 60 mm diameter). Eggs for experimental flies were collected from these ‘standardised’ population cages.

### 2.6 Effect of standard and poor diet on post-infection survival

This experiment tested the effect of poor diet on the post-infection survival of the flies, when compared to the standard food, for both EPN and IUS selection regimes. For each experimental population, standardized fly cages were provided with *ad libitum* yeast paste smeared on the top of the banana-jaggery-yeast food plate. After two days, these plates were replaced with oviposition food plates for 18 hours. From these oviposition plates, eggs were collected and distributed randomly into 20 vials containing standard diet (100% of standard food composition, 6-8 ml per vial) and 20 vials containing poor diet (50% diluted standard food; every component of the standard food composition was reduced to half of the original except water, agar, and preservatives; 6-8 ml per vial) at a density of 60-80 eggs per vial. These vials were incubated under standard laboratory conditions for 12 days PEL. Peak eclosion happens on 10^th^ day PEL and by 12^th^ day PEL, flies would have matured and mated at least once in the rearing vial itself. Please note that the eclosing adults stayed in the rearing vials till the day of infection and hence continued to be on same standard or poor diet in which they were reared as larvae.

On day 12 PEL, flies from each population, reared on either standard or poor diet, were randomly assigned to one of the following treatments: (a) infected with pathogen: 100 females and 100 males divided into two cages with equal density and sex ratio; and (b) sham-infected: 100 females and 100 males divided into two cages with equal density and sex ratio. Post-treatment the flies were housed in plexiglass cages (14 cm x 16 cm x 13 cm) provided with ad libitum access to either standard or poor diet (depending on the diet they were raised in as larvae). Hence, flies remained on the same diet throughout their life: as larvae, before infection, and after infection. Mortality of the flies were recorded every 4-6 hours for 96 hours after infection. Therefore, total 96 cages [2 cages × 2 treatments (infected or sham) × 2 diet × 3 populations × 4 blocks] was observed for each selection regime.

This experiment was carried out using the EPN selection regime after 40 generations of forward selection, and with the IUS selection regime after 145 generations of forward selection. Additionally, flies from EPN selection regime were infected with *E. faecalis* (infection dose: OD_600_ = 1.0) and flies from IUS selection regime were infected with *P. entomophila* (infection dose: OD_600_ = 1.5). For logistic ease, the experiment was carried out one block on each day, i.e., E1, P1, and N1 (or I1, U1, and S1) were handled together of one day, and so on.

### 2.7 Effect of standard and poor diet on female fecundity

Along with the assay for differences in post-infection survival, we assayed for the effect of diet, infection status, and selection history on female fecundity. 96 hours after infection (or, sham-infection), the above fly cages were provided with oviposition food-plates for the flies to lay eggs on for 18 hours. After 18 hours, these plates were withdrawn, labelled and stored at −20°C and eggs were counted later. Per-female fecundity was calculated by dividing the number of eggs layed during the 18-hour window by the number of females alive in that cage at the start of the oviposition period. The oviposition food plates were of the same diet (standard or poor) the flies were being held on till that point.

### 2.8 Effect of standard and poor diet on egg-to-adult development time and viability

In this experiment we tested if rearing on standard vs. poor diet affected the egg-to-adult development time and viability of flies from both EPN and IUS selection regime.

Two days prior to the egg collection, fresh food plates (normal food composition), smeared with yeast paste, were provided to the standardized fly cages. On the day of egg collection, similarly yeasted food-plate was provided for 6 hours and withdrawn. This was followed by a second and a third round of yeasted food-plate, each for an hour only. This was done to encourage the females to lay the stored eggs. Following these, a fresh food plate was provided to the cages for 1 hour, and eggs were collected from this plate to start the assay.

From each population, 20 vials with exactly 70 eggs each were set up: 10 with standard diet and 10 with poor diet. These vials were incubated under standard laboratory conditions. Once flies started eclosing, flies were transferred into fresh empty vials, every 4 hours, and labelled according to vial, population, and diet identity. This was done until the very last fly eclosed. These freshly eclosed flies were immediately frozen in −20°C, and later sexed and counted.

This experiment was carried out using the EPN selection regime after 38 generations of forward selection, and with the IUS selection regime after 85 generations of forward selection. For logistic ease, the experiment was carried out one block on each day, i.e., E1, P1, and N1 (or I1, U1, and S1) were handled together of one day, and so on.

### 2.9 Effect of standard and poor diet on dry body weight (at eclosion)

Measurement of dry body weight at eclosion was done using flies stored at the end of the development time assay. For each population of the IUS selection regime, flies eclosing from the same vial were pooled together according to vial identity. For each population within each diet treatment, 5 flies of each sex were randomly picked from the pooled sample, and transferred to 1.5 ml micro-centrifuge tubes. The sexes were kept in separate tubes. Therefore, each vial from the development time assay produced one tube of each sex. These tubes were dry heated in hot air oven for 48 hours at 60°C before being weighed. Weight was measured using Sartorius weighing balance (model CPA225D). Dry body weight of the flies of EPN populations were not measured as there was no difference in the development time.

### 2.10 Statistical analysis

All statistical analysis was done in R statistical software (version 4.1, R Core Team 2021). Post-infection survival of flies was modelled as

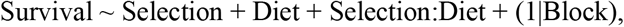

using mixed-effect Cox proportional hazards model (*coxme* function from ‘coxme’ package, Therneau 2020). Test for significant effects of different factors in the model was carried out using analysis of deviance (*Anova* function from ‘car’ package). Data from each sex was analyzed separately.

Life-history traits were analyzed using mixed-effect general linear models (*lmer* function from ‘lmerTest’ package) and subjected to type III ANOVA (*anova* function from base R) for significance tests. The mixed-effect linear models used were as follows:

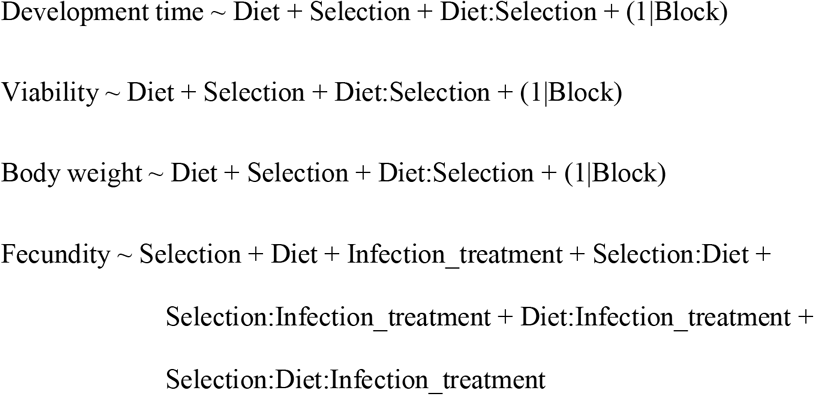

## 3. RESULTS

### 3.1 Effect of poor larval diet on immune function and life-history traits of IUS selection regime

Flies from I (selected against *Pseudomonas entomophila*), S (sham-infected controls), and U (uninfected controls) populations were raised as larvae on two different diets: standard diet and poor diet (50 percent reduction in nutritious diet components). Adult flies were hosted on the same diet as larva during the course of the experiment. We tested the effect of rearing on different diets on (a) immune function and (b) fecundity of adult flies, (c) egg-to-adult development time and survival, and (d) adult dry body weight.

To test for the effect of diet on immune function, adult flies of both sexes from all populations were infected with *Pseudomonas entomophila*, along with sham-infected controls, and their mortality was recorded for 96 hours post-infection. There was negligible mortality in sham-infected flies of all populations (Figures 2A and 2B), hence data from only the infected flies was analyzed for effect of selection, diet, and selection × diet interaction; sexes were analyzed separately. Selection history had a significant effect on post-infection survival in females (Figure 2A, χ^2^_(df=2)_ = 383.1930, p < 2.2e-16): I females survived better than S and U females. There was a significant effect of selection × diet interaction (Figure 2A, χ^2^_(df=2)_ = 6.2361, p = 0.04424): U and S females raised on poor diet survived less compared to flies raised on standard diet; I females survived equally well irrespective of the diet they were raised on. Selection history also had a significant effect on post-infection survival in males (Figure 2B, χ^2^_(df=2)_ = 441.6823, p < 2.2e-16): I males survived better than S and U males. Males raised on poor larval diet were more susceptible to infection compared to males raised on standard diet (χ^2^_(df=1)_ = 7.8464, p = 0.005092), but no significant effect of selection × diet (χ^2^_(df=2)_ = 4.6308, p = 0.098727) was observed on survival of infected males (Figure 2B).

**Figure 2.**
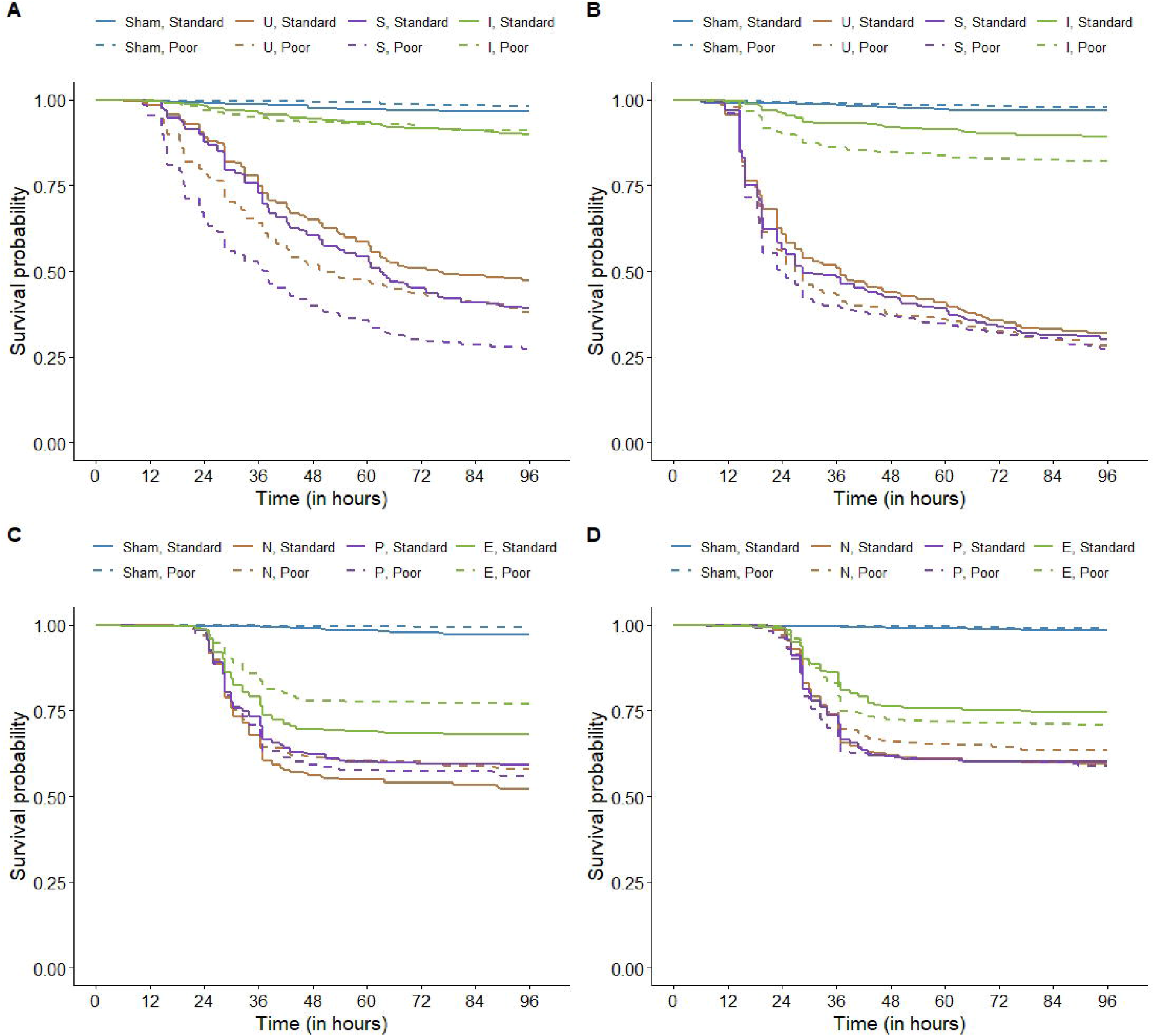
Effect of standard vs. poor larval diet on immune function (post-infection survival) of (a) females and (b) males from I, U, and S populations (infected with *Pseudomonas entomophila*), and (c) females and (d) males from E, P, and N populations (infected with *Enterococcus faecalis*).

Along with immune function assay, we tested for differences in fecundity of flies of all three populations, raised on both diets, when infected or sham-infected (see METHODS, section 2.7, for more details). Diet had a significant effect on female fecundity (F_1,92_ = 17.6550, p = 6.122e-05): females raised on poor diet had less per-capita fecundity compared to females raised on standard diet. Selection history and infection status (and interaction between these and with diet) had no effect on female fecundity (Table 2A, Figure 3A). Block (random factor) had no significant effect on female fecundity (log-likelihood = −303.29, p = 0.2186).

**Table 1.** Analysis of deviance on mixed-effect Cox proportional hazards models for effect of selection history and larval diet on post-infection survival of flies (blocks used as random factors).

| Terms of model | $\chi^2$ | df | p |
| --- | --- | --- | --- |
| (A) Survival of females from I, U, S populations infected with *Pseudomonas entomophila* |  |  |  |
| Selection | 383.1930 | 2 | < 2.2 e-16 |
| Diet | 34.5336 | 1 | 4.19 e-09 |
| Selection:Diet | 6.2361 | 2 | 0.04424 |
| (B) Survival of males from I, U, S populations infected with *Pseudomonas entomophila* |  |  |  |
| Selection | 441.6823 | 2 | < 2.2 e-16 |
| Diet | 7.8464 | 1 | 0.005092 |
| Selection:Diet | 4.6308 | 2 | 0.098727 |
| (C) Survival of females from E, P, N populations infected with *Enterococcus faecalis* |  |  |  |
| Selection | 56.2652 | 2 | 6.056 e-13 |
| Diet | 3.0068 | 1 | 0.08291 |
| Selection:Diet | 8.2023 | 2 | 0.01655 |
| (D) Survival of males from E, P, N populations infected with *Enterococcus faecalis* |  |  |  |
| Selection | 37.1255 | 2 | 8.676 e-09 |
| Diet | 0.1018 | 1 | 0.7496 |
| Selection:Diet | 2.6123 | 2 | 0.2709 |

**Table 2.**
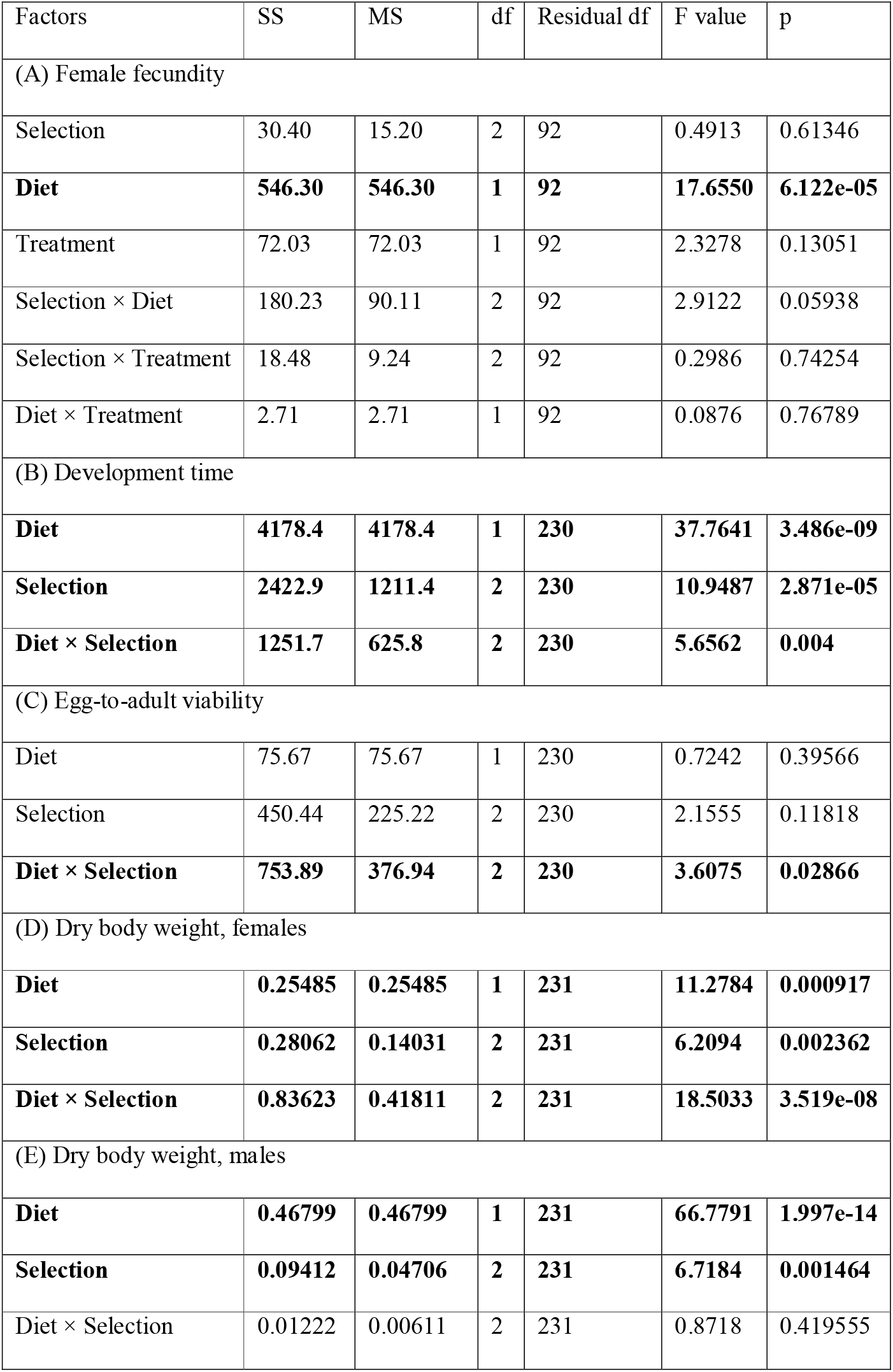
Analysis of variance (type III sum of squares) for the effect of selection history and larval diet (and infection status, if applicable) on life-history traits of flies from I, U, and S populations (blocks used as random factors). Significant effects are marked in bold letters.

**Figure 3.**
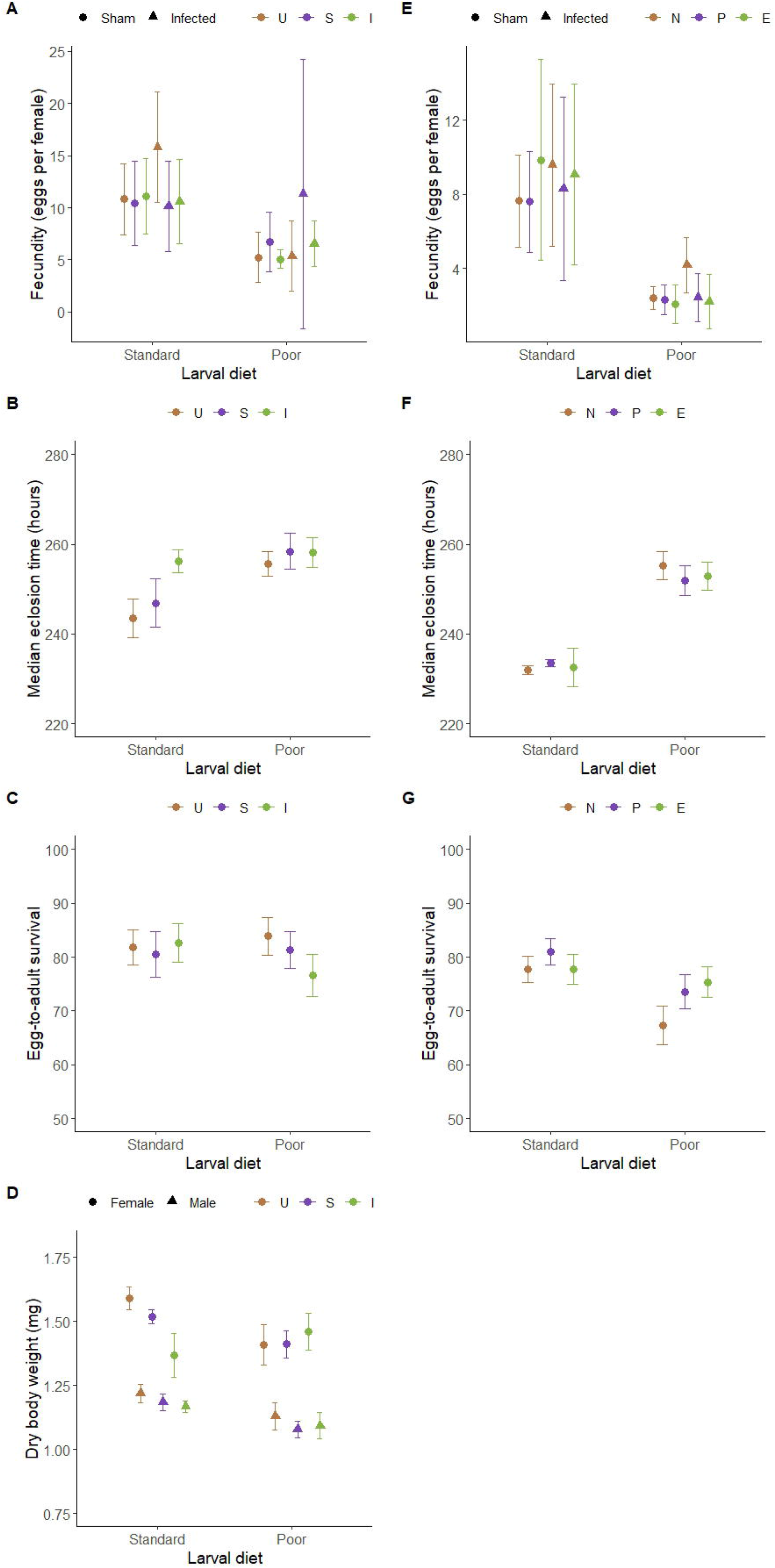
Effect of standard vs. poor larval diet on fecundity (a, e), egg-to-adult development time (b, f) and survival (c, g), and dry body weight at eclosion (d), of flies from I, U, and S populations (a-d) and E, P, and N populations (e-g).

Egg-to-adult development time of flies reared on poor diet was significantly longer than flies reared on standard diet (F_1,230_ = 37.7641, p = 3.486e-09). There was also a significant effect of diet × selection history interaction on development time (F_2,230_ = 5.6562, p = 0.004, Figure 3B): among flies reared in standard diet, those from I populations took significantly longer to develop compared to both S (Tukey’s HSD, p = 0.0021) and U (Tukey’s HSD, p < 0.0001) flies. No such effect of selection history was observed among flies reared on poor diet. Egg-to-adult survival was also affected by diet × selection history interaction (Table 2C, Figure 3C): among flies reared in poor diet, survival was reduced in case of I flies compared to U flies (Tukey’s HSD, p = 0.0199). Block (random factor) had significant effect on both development time and egg-to-adult survival (development time: log-likelihood = −904.39, p = 8445e-09; survival: log-likelihood = −895.83, p = 4.674e-08).

Dry body weight at eclosion for females was significantly affected by larval diet, selection history and the interaction between the two (Table 2D, Figure 3D). Females reared on poor diet overall had lower body weight at eclosion (F_1,231_ = 11.2784, p = 0.000917). Within the females reared on standard diet, I females had lower body weight compared to both U (Tukey’s HSD, p < 0.0001) and S females (Tukey’s HSD, p = 0.0002). No such effect of selection history was apparent among females raised on poor diet. In fact, I females raised on standard diet did not differ in body weight from I females raised on poor diet (Tukey’s HSD, p = 0.0536). Dry body weight at eclosion for males was significantly affected by larval diet and selection history, but not their interaction (Table 2E, Figure 3D). Males reared on poor diet had lower body weight at eclosion (F_1,231_ = 66.7791, p = 1.997e-14). Across both diets, males of I and S populations had lower body weight compared to those of U population (Tukey’s HSD, p = 0.0055 and 0.0056 respectively); weights of males of I and S populations were not different from one another (Tukey’s HSD, p = 0.9997). Block (random factor) had significant effect on dry body weight of both females and males (females: log-likelihood = 48.817, p = <2.2e-16; males: log-likelihood = 165.20, p = <2.2e-16).

### 3.2 Effect of poor larval diet on immune function and life-history traits of EPN selection regime

Flies from E (selected against *Enterococcus faecalis*), P (sham-infected controls), and N (uninfected controls) populations were raised as larvae on two different diets: standard diet and poor diet (50 percent reduction in nutritious diet components). Adult flies were hosted on the same diet as larva during the course of the experiment. We tested the effect of rearing on different diets on (a) immune function and (b) fecundity of adult flies, and (c) egg-to-adult development time and survival.

To test for effect of diet on immune function, adult flies of both sexes from all populations were infected with *Enterococcus faecalis*, along with sham-infected controls, and their mortality was recorded for 96 hours post-infection. There was negligible mortality in sham-infected flies of all populations (Figures 2C and 2D), hence data from only the infected flies was analyzed for effect of selection, diet, and selection × diet interaction; sexes were analyzed separately. Selection history had a significant effect on post-infection survival in females (Figure 2C, χ^2^_(df=2)_ = 56.2652, p = 6.056e-13): E females survived better than P and N females. Overall, across all populations, larval diet had no effect on post-infection survival of females (Figure 2C, χ^2^_(df=1)_ = 3.0068, p = 0.08291). There was a significant effect of selection × diet interaction (Figure 2C, χ^2^_(df=2)_ = 8.2023, p = 0.01655): E females raised on poor diet were less susceptible to infection compared to E females raised on standard diet; no such difference was observed in case of P and N females. Selection history also had a significant effect on post-infection survival in males (Figure 2D, χ^2^_(df=2)_ = 37.1255, p = 8.767e-09): E males survived better than P and N females. No significant effect of diet (χ^2^_(df=1)_ = 0.1018, p = 0.7496) or selection × diet (χ^2^_(df=2)_ =2.6123, p = 0.2709) was observed on survival of infected males (Figure 2D).

Diet had a significant effect on female fecundity (F_1,92_ = 104.4017, p < 2e-16): females raised on poor diet had less per-capita fecundity compared to females raised on standard diet across all three populations. Selection history and infection status (and interaction between these and with diet) had no effect on female fecundity (Table 3A, Figure 3E). Block (random factor) had no significant effect on female fecundity (log-likelihood = −256.88, p = 3.015e-07).

**Table 3.** Analysis of variance (type III sum of squares) for the effect of selection history and larval diet (and infection status, if applicable) on life-history traits of flies from E, P, and N populations (blocks used as random factors). Significant effects are marked in bold letters.

| Factors | SS | MS | df | Residual df | F value | p |
| --- | --- | --- | --- | --- | --- | --- |
| (A) Female fecundity: |  |  |  |  |  |  |
| Selection | 11.30 | 5.65 | 2 | 92 | 0.6669 | 0.5157 |
| **Diet** | **884.60** | **884.60** | **1** | **92** | **104.4017** | **<2e-16** |
| Treatment | 10.08 | 10.08 | 1 | 92 | 1.1897 | 0.2782 |
| Selection × Diet | 18.71 | 9.36 | 2 | 92 | 1.1042 | 0.3358 |
| Selection × Treatment | 19.42 | 9.71 | 2 | 92 | 1.1461 | 0.3224 |
| Diet × Treatment | 0.02 | 0.02 | 1 | 92 | 0.0019 | 0.9651 |
| (b) Development time |  |  |  |  |  |  |
| **Diet** | **25295.0** | **25295.0** | **1** | **231.02** | **373.7692** | **<2e-16** |
| Selection | 39.3 | 19.6 | 2 | 231.02 | 0.2901 | 0.7485 |
| Diet × Selection | 254.3 | 127.2 | 2 | 231.02 | 1.8789 | 0.1551 |
| (c) Egg-to-adult viability |  |  |  |  |  |  |
| **Diet** | **2659.72** | **2659.72** | **1** | **230.98** | **39.1141** | **1.916e-09** |
| **Selection** | **1005.41** | **502.70** | **2** | **230.98** | **7.3928** | **0.0007725** |
| **Diet × Selection** | **649.59** | **324.79** | **2** | **230.98** | **4.7764** | **0.0092762** |

Only larval diet had a significant effect on egg-to-adult development time (Table 3B, Figure 3F), with all three populations taking longer to develop when reared in poor diet (F_1,231_ = 373.7692, p < 2e-16). Flies reared on poor larval diet also had reduced egg-to-adult survival (F_1,231_ = 39.1141, p = 1.916 e-09, Figure 3G). Egg-to-adult survival was also significantly affected by selection history, and selection × diet interaction (Table 3C, Figure 3G): within flies reared on poor diet, flies of N population had reduced survival compared both P (Tukey’s HSD, p = 0.0133) and E (Tukey’s HSD, p = 0.0.004) population, which did not differ among themselves (Tukey’s HSD, p = 0.9327). No such effect of selection was apparent within flies reared on standard diet. Block (random factor) had significant effect on both development time and egg-to-adult survival (development time: log-likelihood = −845.53, p = 7.307e-07; survival: log-likelihood = −845.87, p = 9.72e-07).

## 4. DISCUSSION

### 4.1 Adult immune function

Selection history had a dominating influence on adult immune function. Flies from populations experimentally selected for improved immune function (I and E) exhibited significantly less mortality when infected, compared to their corresponding controls (U and S, and P and N populations, respectively), irrespective of the quality of the diet the flies were reared on (Figure 2). Improvement of immune function, in response to experimental evolution, therefore, is not lost due to resource scarcity.

Rearing on poor larval diet reduces post-infection survival of flies in a pathogen specific manner. When flies of IUS selection regime are infected with *Pseudomonas entomophila* we see a marked decrease in survival of flies reared on poor larval diet (Figures 2A and 2B). On the other hand, when EPN flies are infected with *Enterococcus faecalis* survival of flies was not affected by whether they were raised on standard or poor diet (Figures 2C and 2D). These results agree with that of previous studies that have shown that host’s access to nutrition determines their resistance against *P. entomophila* (Kutzer et al. 2017), but not against *E. faecalis* (Ayres and Schneider 2009). There can be a few possible reasons for this observation. First, defense against *E. faecalis* may be less resource intensive compared to defense against *P. entomophila*. Flies with chronic *E. faecalis* infection do not exhibit any change in starvation resistance, while such change is observable for chronic infection with other pathogens (Chambers et al. 2019). Flies housed at high densities, where resources become limiting, show little or no reduction of defense against *E. faecalis*, while reduction in defense against other pathogens is clearly apparent (Das et al. 2021, preprint). Second, mechanisms utilized by flies to defend against the two pathogens may be different (Lemaitre and Hoffman 2007), and different mechanisms are known to be affected by resource limitation to different extents (Adamo et al. 2016). In this study, we did not test for the effect of poor diet on defense against *E. faecalis* of I flies (selected for defense against *P. entomophila*) or defense against *P. entomophila* of E flies (selected for defense against *E. faecalis*). Therefore, we are unable to comment on any role of selection history × pathogen identity in determining the effect of poor diet on immune function. Our previous results have shown that the I flies (compared to S controls) exhibit cross-resistance against *E. faecalis* and E flies (compared to P controls) exhibit cross-resistance against *P. entomophila* (Singh et al. 2021, preprint). Therefore, it will be interesting to explore this avenue further in future experiments.

The effect of poor diet on immune function of the host is dependent upon both the selection history of the host and the sex of the host. Males from I, U, and S populations exhibited greater mortality when reared on poor larval diet, but such increase in mortality was only observable in females of U and S populations (Figures 2A and 2B); females of I populations survived equally well when infected irrespective of the larval diet they were reared on. On the other hand, males from E, P, and N populations, and females from P and N populations, do not show any larval diet dependent difference in post-infection survival, but E females exhibit decreased mortality when raised on poor diet (Figures 2C and 2D). Therefore, counterintuitively, females of both the selected populations (I and E) exhibit reduced post-infection mortality when raised in poor diet than what would be expected of them if the response of their corresponding controls are considered typical for flies. Sex-specific differential effect of diet quality is also observed in field cricket, *Gryllus texensis*, where high quality diet increases survival of males, but compromises survival of females, when the crickets are infected with *Serratia marcescens* (Kelly and Tawes 2013). Although sexual dimorphism in immune function can stem from various sources (Zuk and McKean 1996, Rolf et al. 2002, Nunn et al. 2009, Vincent and Sharp 2014, Sharp and Vincent 2015, Belmonte et al. 2020), the reason for sex-specific diet × selection history interaction is not very obvious. Resource allocation priorities can shift depending upon environmental factors and resource levels (Ng’oma et al. 2017), and it is possible that adaptation to regular bacterial challenge involves prioritizing investment towards immune function, especially when resources are scarce. Females of the selected populations, therefore, may have evolved to prioritize immune defense over other function, compared to corresponding controls, especially when resources are scarce. Reduced fecundity in females due to poor quality diet may also free up whatever limited resources, which can then be channelized towards immune defense (see section 4.2). Excess resources in form of dietary yeast supplement are known to ameliorate reproduction-immunity trade-off in females, but not in male *Drosophila melanogaster* (McKean and Nunney 2005). Not much is known about the effect of larval malnutrition of male reproductive capacity in flies.

Another reason why any population – E, P, or N – when reared on poor larval diet do not become more susceptible to infection with *E. faecalis* may be due to reduced insulin signalling. Poor larval diet is known to reduce insulin signalling in flies (Rehman and Vargheshe 2021), and inactivated insulin signalling makes flies more resistant to infection with *E. faecalis* (Libert et al 2008). This might explain why in E females we see an increase in post-infection survival when reared on poor larval diet: a combination of evolved increased investment into immune function, reduction in insulin signalling, and reduced investment towards reproduction (later two influenced by poor larval diet) may drive this phenomenon.

### 4.2 Fecundity

In both selection regimes, IUS and EPN, female fecundity was only affected by larval diet, with flies from poor diet producing less eggs than flies from standard diet (Figures 3A and 3E). Larval access to nutrition is a major determinant of adult reproductive fitness across many different Dipteran species (Hodin 2009). Poor larval diet is known to reduce adult fecundity in female *Drosophila melanogaster*, either directly due to reduced resources or indirectly due to reduction in both ovariole number and body size (Hodin and Riddiford 2000, Tu and Tatar 2003, Deas et al. 2019, Klepsatel et al. 2020). Decreased ovariole count also implies a smaller *resource sink*: less ovarioles equals to less opportunity to allocate resources towards reproduction. This can potentially free up resources, that would otherwise have been invested in oogenesis, and even in times of resource scarcity, help maintain other organismal functions (such as immune function) at their optimal levels. Ablation of germline, and therefore the *resource sink*, has been previously shown to improve immune function in flies (Short et al. 2012, Rodrigues et al. 2021)

Neither infection status, nor selection history had any effect on female fecundity in our experiments (Figures 2A and 2E). Flies experimentally evolved to defend against *P. entomophila* have previously been shown to not pay any fecundity cost of improved resistance (Faria et al. 2015, Gupta et al. 2015). Here we confirm the results for *P. entomophila*, and also report absence of fecundity costs when flies adapt to defend against *E. faecalis*. Reduced fecundity, either because of increased genetic resistance to pathogens or because of the energetic cost of mounting an immune response, is a typical predicted manifestation of cost of immunity (Lochmiller and Deerenberg 2000, Schmid-Hempel 2003, McKean et al. 2008). Despite of this assertion, reduction of fecundity in response to infection is not always observed in experiments with *D. melanogaster*. For example, infection with *P. entomophila, L. lactis*, and *E. coli* does not lead to fecundity decline in female flies (Kutzer and Armitage 2016, Kutzer et al. 2018), while infection with *P. aeruginosa* is even known to increase fecundity (Hudson et al. 2020, but see Linder and Promislow 2009). Various factors, like pathogen identity, host genotype, infection route, and whether the bacteria colonize the ovaries, may potentially underlie the observed variation in experimental outcome (Brandt and Schneider 2007, Linder and Promislow 2009, Gupta et al. 2017). An additional determining factor can be the time of fecundity measurement relative to the time of infection. In our study, we measured fecundity after the period of infection-induced acute mortality had passed. Differences in fecundity due to cost of immune activation may be more apparent if measured during the acute phase of infection.

### 4.3 Egg-to-adult development time and viability

In both selection regimes, IUS and EPN, rearing on poor larval diet negatively affected various larval traits. In IUS, rearing on poor larval diet led to increase in development time and reduction in dry body weight at eclosion, although egg-to-adult viability was unaffected (Figure 3B, 3C, 3D). In EPN, poor larval diet increased development time and reduced egg-to-adult viability (Figure 3F and 3G). Multiple previous studies have demonstrated similar effects of poor larval diet on larval traits (Kolss et al 2009, Deas et al 2019), with the effects being primarily driven by reduction in protein content in the diet (Tu and Tatar 2003, Klepsatel et al 2020). Interestingly enough, poor or low protein larval diet increases adult lipid reserves, and increases starvation resistance by slowing down metabolic rate, without any effect on adult lifespan (Tu and Tatar 2003, Klepsatel et al 2019, Rehman and Varghese 2021).

## 5. CONCLUSION

As discussed above (see section 4), developing as larvae in resource poor environments can affect numerous adult traits in *Drosophila melanogaster*, including body size and reproductive capacity. Our results show that populations experimentally selected to defend against pathogen challenge can become better at counteracting the effect of poor larval nutrition on adult immune function. For example, poor larval diet had no effect on post-infection survival of I population females (selected for defense against *P. entomophila*), while the corresponding control U and S population females showed reduced post-infection survival upon being reared on poor larval diet (Figure 2A). Interestingly, the selected populations exhibited similar depression of female fecundity, induced by poor larval diet, as exhibited by the control population (Figure 3A and 3E). This indicates that since adult immune function is under direct selection in the I populations, these flies have evolved to prioritize investment towards immune function even under circumstances where resources are limited. We see no such selection history-dependent differential effect of larval diet in E population females (selected for defense against *E. faecalis*), and the corresponding control P and N population females (Figure 2C), because larval diet does not affect immune defense against *E. faecalis*, which is the selective agent in this case.

Adaptation to poor larval diet has been shown to increase susceptibility of selected *Drosophila melanogaster* population to oral infection (Vijendravarma et al 2015); this increase in susceptibility is not driven by resource scarcity, but is due to the evolution of increased gut permeability as an adaptation to malnutrition. These selected populations are also better at counteracting the negative effects of poor larval diet on various other life-history traits (Kolss et al 2009). It would be interesting to test how these populations fair in terms of adult immune function (systemic pathogen challenge) given that immune function is not directly under selection in these populations.

Previous theory and empirical research have suggested that life-history trade-offs associated with increased immune function are expected to be more overt under low resource environment (Lazzaro and Little 2009). Results from our experiments do not agree with this expectation. In our experiments, although most measured life-history traits were negatively affected by poor larval diet, the selected populations were not adversely affected compared to the control populations when reared on poor diet (Figure 3). In fact, I population flies exhibited prolonged development time and reduced dry body weight at eclosion (only for females) compared to controls (U and S population flies) when reared on standard diet, but not when reared on poor larval diet. Hence trade-offs were observed when resources were abundant, and not when resources were limited. We did not observe any trade-offs in the E, P, and N populations on either diet.

To summarize, in this study we explored if poor larval nutrition has an effect on adult immunity and other life-history traits in *Drosophila melanogaster* populations experimentally evolved to be immune to bacterial infection. Our results suggest that (a) effect of poor larval nutrition on adult defense against bacterial infection is pathogen specific; (b) experimentally evolved populations maintain a better functioning immune system, compared to control populations, even when raised on poor diet; (c) host sex and selection history interact to determine the effect of poor diet on adult immune function; (d) poor larval diet reduces females fecundity, but fecundity is not affected by either host selection history or infections status; (e) poor larval diet prolongs egg-to-adult development time; and, (f) cost of evolved immune defense can manifest in form of prolonged egg-to-adult development, depending upon the pathogen used for selection. We therefore conclude that effect of poor nutrition on host immune function is not uniform, but contingent upon host sex, level of host’s resistance to pathogen (selection history), and very importantly, the identity of the pathogen.

## Acknowledgements

This study was funded by IISER Mohali intramural funds and a research grant (no. BT/PR14278/BRB/10/1417/2015) from Dept. of Biotechnology, Govt. of India. AS is supported by Junior and Senior Research Fellowships from University Grant Council, Govt. of India. AKB is supported by Junior and Senior Research Fellowships from Council of Scientific and Industrial Research, Govt. of India. BS is supported by INSPIRE Scholarship for Higher Education, and TH and NB are supported by KVPY Fellowship, both from Dept. of Science and Technology, Govt. of India. We thank Prof. P Cornelis (Free University, Brussels, Belgium) for providing us *Pseudomonas entomophila*, Prof. B Lazzaro (Cornell University, Ithaca, USA) for providing us *Enterococcus faecalis*. IUS selection regimes used in this study was originally established by Dr. V Gupta during her Ph.D. in Prof. N G Prasad’s laboratory. We would like to thank Nagender Kumar for all the help during experiments.

## Notes

Conflict of interests: Authors declare no competing interests.

### Competing Interest Statement

The authors have declared no competing interest.

## REFERENCES

1. Abbott, J., 2014. Self-medication in insects: current evidence and future perspectives. Ecological Entomology 39, 273–280. https://doi.org/10.1111/een.12110

2. Adamo, S.A., Davies, G., Easy, R., Kovalko, I., Turnbull, K.F., 2016. Reconfiguration of the immune system network during food limitation in the caterpillar Manduca sexta. Journal of Experimental Biology jeb.132936. https://doi.org/10.1242/jeb.132936

3. Ayres, J.S., Schneider, D.S., 2009. The Role of Anorexia in Resistance and Tolerance to Infections in Drosophila. PLoS Biol 7, e1000150. https://doi.org/10.1371/journal.pbio.1000150

4. Bashir-Tanoli, S., Tinsley, M.C., 2014. Immune response costs are associated with changes in resource acquisition and not resource reallocation. Funct Ecol 28, 1011–1019. https://doi.org/10.1111/1365-2435.12236

5. Bedhomme, S., Agnew, P., Sidobre, C., Michalakis, Y., 2004. Virulence reaction norms across a food gradient. Proc. R. Soc. Lond. B 271, 739–744. https://doi.org/10.1098/rspb.2003.2657

6. Belmonte, R.L., Corbally, M.-K., Duneau, D.F., Regan, J.C., 2020. Sexual Dimorphisms in Innate Immunity and Responses to Infection in Drosophila melanogaster. Frontiers in Immunology 10.

7. Boots, M., Begon, M., 1993. Trade-Offs with Resistance to a Granulosis Virus in the Indian Meal Moth, Examined by a Laboratory Evolution Experiment. Functional Ecology 7, 528–534. https://doi.org/10.2307/2390128

8. Brandt, S.M., Schneider, D.S., 2007. Bacterial infection of fly ovaries reduces egg production and induces local hemocyte activation. Developmental & Comparative Immunology 31, 1121–1130. https://doi.org/10.1016/j.dci.2007.02.003

9. Chambers, M.C., Jacobson, E., Khalil, S., Lazzaro, B.P., 2019. Consequences of chronic bacterial infection in Drosophila melanogaster. PLOS ONE 14, e0224440. https://doi.org/10.1371/journal.pone.0224440

10. Cotter, S.C., Simpson, S.J., Raubenheimer, D., Wilson, K., 2011. Macronutrient balance mediates trade offs between immune function and life history traits. Functional Ecology 25, 186–198. https://doi.org/10.1111/j.1365-2435.2010.01766.x

11. Cressler, C.E., Nelson, W.A., Day, T., McCauley, E., 2014. Disentangling the interaction among host resources, the immune system and pathogens. Ecology Letters 17, 284–293. https://doi.org/10.1111/ele.12229

12. Das, P.N., Basu, A., Prasad, N.G., 2022. Increasing adult density compromises anti-bacterial defense in Drosophila melanogaster (preprint). Evolutionary Biology. https://doi.org/10.1101/2022.01.02.474745

13. Deas, J.B., Blondel, L., Extavour, C.G., 2019. Ancestral and offspring nutrition interact to affect life-history traits in Drosophila melanogaster. Proceedings of the Royal Society B: Biological Sciences 286, 20182778. https://doi.org/10.1098/rspb.2018.2778

14. Diamond, S.E., Kingsolver, J.G., 2011. Host plant quality, selection history and trade-offs shape the immune responses of *Manduca sexta*. Proc. R. Soc. B. 278, 289–297. https://doi.org/10.1098/rspb.2010.1137

15. Dinh, H., Mendez, V., Tabrizi, S.T., Ponton, F., 2019. Macronutrients and infection in fruit flies. Insect Biochemistry and Molecular Biology 110, 98–104. https://doi.org/10.1016/j.ibmb.2019.05.002

16. Faria, V.G., Martins, N.E., Paulo, T., Teixeira, L., Sucena, É., Magalhães, S., 2015. Evolution of Drosophila resistance against different pathogens and infection routes entails no detectable maintenance costs: EVOLUTION OF RESISTANCE HAS NO MAINTENANCE COSTS. Evolution 69, 2799–2809. https://doi.org/10.1111/evo.12782

17. Fellowes, M.D.E., Kraaijeveld, A.R., Godfray, H.C.J., 1998. Trade–off associated with selection for increased ability to resist parasitoid attack in Drosophila melanogaster. Proc. R. Soc. Lond. B 265, 1553–1558. https://doi.org/10.1098/rspb.1998.0471

18. Gupta, V., Venkatesan, S., Chatterjee, M., Syed, Z.A., Nivsarkar, V., Prasad, N.G., 2016. No apparent cost of evolved immune response in Drosophila melanogaster. Evolution 70, 934–943. https://doi.org/10.1111/evo.12896

19. Gupta, V., Vasanthakrishnan, R.B., Siva-Jothy, J., Monteith, K.M., Brown, S.P., Vale, P.F., 2017. The route of infection determines Wolbachia antibacterial protection in Drosophila. Proceedings of the Royal Society B: Biological Sciences 284, 20170809. https://doi.org/10.1098/rspb.2017.0809

20. Harshman, L.G., Hoffmann, A.A., 2000. Laboratory selection experiments using Drosophila: what do they really tell us? Trends in Ecology & Evolution 15, 32–36. https://doi.org/10.1016/S0169-5347(99)01756-5

21. Hite, J.L., Pfenning, A.C., Cressler, C.E., 2020. Starving the Enemy? Feeding Behavior Shapes Host-Parasite Interactions. Trends in Ecology & Evolution 35, 68–80. https://doi.org/10.1016/j.tree.2019.08.004

22. Hodin, J., 2009. She Shapes Events as they Come: Plasticity in Female Insect Reproduction, in: Whitman, D., Ananthakrishnan, T. (Eds.), Phenotypic Plasticity of Insects. Science Publishers. https://doi.org/10.1201/b10201-12

23. Hodin, J., Riddiford, L.M., 2000. Different Mechanisms Underlie Phenotypic Plasticity and Interspecific Variation for a Reproductive Character in Drosophilids (insecta: Diptera). Evolution 54, 1638–1653. https://doi.org/10.1111/j.0014-3820.2000.tb00708.x

24. Hudson, A.L., Moatt, J.P., Vale, P.F., 2020. Terminal investment strategies following infection are dependent on diet. Journal of Evolutionary Biology 33, 309–317. https://doi.org/10.1111/jeb.13566

25. Kelly, C.D., Tawes, B.R., 2013. Sex-Specific Effect of Juvenile Diet on Adult Disease Resistance in a Field Cricket. PLoS ONE 8, e61301. https://doi.org/10.1371/journal.pone.0061301

26. Klepsatel, P., Procházka, E., Gáliková, M., 2018. Crowding of Drosophila larvae affects lifespan and other life-history traits via reduced availability of dietary yeast. Experimental Gerontology 110, 298–308. https://doi.org/10.1016/j.exger.2018.06.016

27. Klepsatel, P., Knoblochová, D., Girish, T.N., Dircksen, H., Gáliková, M., 2020. The influence of developmental diet on reproduction and metabolism in Drosophila. BMC Evol Biol 20, 93. https://doi.org/10.1186/s12862-020-01663-y

28. Kolss, M., Vijendravarma, R.K., Schwaller, G., Kawecki, T.J., 2009. LIFE-HISTORY CONSEQUENCES OF ADAPTATION TO LARVAL NUTRITIONAL STRESS IN DROSOPHILA. Evolution 63, 2389–2401. https://doi.org/10.1111/j.1558-5646.2009.00718.x

29. Kraaijeveld, A.R., Godfray, H.C.J., 1997. Trade-off between parasitoid resistance and larval competitive ability in Drosophila melanogaster. Nature 389, 278–280. https://doi.org/10.1038/38483

30. Kutzer, M.A.M., Armitage, S.A.O., 2016. The effect of diet and time after bacterial infection on fecundity, resistance, and tolerance in Drosophila melanogaster. Ecol Evol 6, 4229–4242. https://doi.org/10.1002/ece3.2185

31. Kutzer, M.A.M., Kurtz, J., Armitage, S.A.O., 2018. Genotype and diet affect resistance, survival, and fecundity but not fecundity tolerance. J. Evol. Biol. 31, 159–171. https://doi.org/10.1111/jeb.13211

32. Lazzaro, B.P., Little, T.J., 2009. Immunity in a variable world. Phil. Trans. R. Soc. B 364, 15–26. https://doi.org/10.1098/rstb.2008.0141

33. Lee, K. p, Cory, J. s, Wilson, K., Raubenheimer, D., Simpson, S. j, 2006. Flexible diet choice offsets protein costs of pathogen resistance in a caterpillar. Proceedings of the Royal Society B: Biological Sciences 273, 823–829. https://doi.org/10.1098/rspb.2005.3385

34. Lemaitre, B., Hoffmann, J., 2007. The Host Defense of Drosophila melanogaster. Annual Review of Immunology 25, 697–743. https://doi.org/10.1146/annurev.immunol.25.022106.141615

35. Libert, S., Chao, Y., Zwiener, J., Pletcher, S.D., 2008. Realized immune response is enhanced in long-lived puc and chico mutants but is unaffected by dietary restriction. Molecular Immunology, Special section: Theories and Modelling of T Cell Behaviour 45, 810–817. https://doi.org/10.1016/j.molimm.2007.06.353

36. Linder, J.E., Promislow, D.E.L., 2009. Cross-generational fitness effects of infection in Drosophila melanogaster. Fly 3, 143–150. https://doi.org/10.4161/fly.8051

37. Lochmiller, R.L., Deerenberg, C., 2000. Trade-offs in evolutionary immunology: just what is the cost of immunity? Oikos 88, 87–98. https://doi.org/10.1034/j.1600-0706.2000.880110.x

38. Marden, J.H., Rogina, B., Montooth, K.L., Helfand, S.L., 2003. Conditional tradeoffs between aging and organismal performance of Indy long-lived mutant flies. PNAS 100, 3369–3373. https://doi.org/10.1073/pnas.0634985100

39. McKean, K.A., Nunney, L., 2005. Bateman’s Principle and Immunity: Phenotypically Plastic Reproductive Strategies Predict Changes in Immunological Sex Differences. Evolution 59, 1510–1517. https://doi.org/10.1111/j.0014-3820.2005.tb01800.x

40. McKean, K.A., Yourth, C.P., Lazzaro, B.P., Clark, A.G., 2008. The evolutionary costs of immunological maintenance and deployment. BMC Evol Biol 8, 76. https://doi.org/10.1186/1471-2148-8-76

41. Miller, C.V.L., Cotter, S.C., 2018. Resistance and tolerance: The role of nutrients on pathogen dynamics and infection outcomes in an insect host. J Anim Ecol 87, 500–510. https://doi.org/10.1111/1365-2656.12763

42. Ng’oma, E., Perinchery, A.M., King, E.G., 2017. How to get the most bang for your buck: the evolution and physiology of nutrition-dependent resource allocation strategies. Proceedings of the Royal Society B: Biological Sciences 284, 20170445. https://doi.org/10.1098/rspb.2017.0445

43. Nunn, C.L., Lindenfors, P., Pursall, E.R., Rolff, J., 2009. On sexual dimorphism in immune function. Philosophical Transactions of the Royal Society B: Biological Sciences 364, 61–69. https://doi.org/10.1098/rstb.2008.0148

44. Pike, V.L., Lythgoe, K.A., King, K.C., 2019. On the diverse and opposing effects of nutrition on pathogen virulence. Proceedings of the Royal Society B: Biological Sciences 286, 20191220. https://doi.org/10.1098/rspb.2019.1220

45. Ponton, F., Lalubin, F., Fromont, C., Wilson, K., Behm, C., Simpson, S.J., 2011. Hosts use altered macronutrient intake to circumvent parasite-induced reduction in fecundity. International Journal for Parasitology 41, 43–50. https://doi.org/10.1016/j.ijpara.2010.06.007

46. Ponton, F., Morimoto, J., Robinson, K., Kumar, S.S., Cotter, S.C., Wilson, K., Simpson, S.J., 2020. Macronutrients modulate survival to infection and immunity in Drosophila. J Anim Ecol 89, 460–470. https://doi.org/10.1111/1365-2656.13126

47. Povey, S., Cotter, S.C., Simpson, S.J., Lee, K.P., Wilson, K., 2009. Can the protein costs of bacterial resistance be offset by altered feeding behaviour? Journal of Animal Ecology 78, 437–446. https://doi.org/10.1111/j.1365-2656.2008.01499.x

48. Prakash, A., Agashe, D., Khan, I., 2022. The costs and benefits of basal infection resistance vs immune priming responses in an insect. Developmental & Comparative Immunology 126, 104261. https://doi.org/10.1016/j.dci.2021.104261

49. R Core Team (2021). R: A language and environment for statistical computing. R Foundation for Statistical Computing, Vienna, Austria. URL: https://www.Rproject.org/.

50. Rehman, N., Varghese, J., 2021. Larval nutrition influences adult fat stores and starvation resistance in Drosophila. PLoS ONE 16, e0247175. https://doi.org/10.1371/journal.pone.0247175

51. Reznick, D., 1985. Costs of Reproduction: An Evaluation of the Empirical Evidence. Oikos 44, 257–267. https://doi.org/10.2307/3544698

52. Rodrigues, M.A., Merckelbach, A., Durmaz, E., Kerdaffrec, E., Flatt, T., 2021. Transcriptomic evidence for a trade-off between germline proliferation and immunity in Drosophila. Evolution Letters 5, 644–656. https://doi.org/10.1002/evl3.261

53. Rolff, J., 2002. Bateman’s principle and immunity. Proceedings of the Royal Society of London. Series B: Biological Sciences 269, 867–872. https://doi.org/10.1098/rspb.2002.1959

54. Rose, M.R., 1984. Laboratory evolution of postponed senescence in Drosophila melanogaster. Evolution, pp.1004–1010. https://doi.org/10.1111/j.1558-5646.1984.tb00370.x

55. Schmid-Hempel, P., 2003. Variation in immune defence as a question of evolutionary ecology. Proceedings of the Royal Society of London. Series B: Biological Sciences 270, 357–366. https://doi.org/10.1098/rspb.2002.2265

56. Sharp, N.P., Vincent, C.M., 2015. The effect of parasites on sex differences in selection. Heredity 114, 367–372. https://doi.org/10.1038/hdy.2014.110

57. Sheldon, B.C., Verhulst, S., 1996. Ecological immunology: costly parasite defences and trade-offs in evolutionary ecology. Trends in Ecology & Evolution 11, 317–321. https://doi.org/10.1016/0169-5347(96)10039-2

58. Short, S.M., Wolfner, M.F., Lazzaro, B.P., 2012. Female Drosophila melanogaster suffer reduced defense against infection due to seminal fluid components. Journal of Insect Physiology 58, 1192–1201. https://doi.org/10.1016/j.jinsphys.2012.06.002

59. Singh, A., Basu, A., Shit, B., Hegde, T., Bansal, N., Prasad, N.G., 2021. Recurrent evolution of cross-resistance in response to selection for improved post-infection survival in Drosophila melanogaster (preprint). Evolutionary Biology. https://doi.org/10.1101/2021.11.26.470139

60. Singh, K., Kochar, E., Prasad, N.G., 2015. Egg Viability, Mating Frequency and Male Mating Ability Evolve in Populations of Drosophila melanogaster Selected for Resistance to Cold Shock. PLOS ONE 10, e0129992. https://doi.org/10.1371/journal.pone.0129992

61. Siva-Jothy, M.T., Thompson, J.J.W., 2002. Short-term nutrient deprivation affects immune function. Physiological Entomology 27, 206–212. https://doi.org/10.1046/j.1365-3032.2002.00286.x

62. Stearns, S.C., 1989. Trade-Offs in Life-History Evolution. Functional Ecology 3, 259–268. https://doi.org/10.2307/2389364

63. Terry M. Therneau (2020). coxme: Mixed Effects Cox Models. R package version 2.2–16. https://CRAN.R-project.org/package=coxme

64. Tu, M.-P., Tatar, M., 2003. Juvenile diet restriction and the aging and reproduction of adult Drosophila melanogaster: Juvenile diet restriction and the aging and reproduction, M.-P. Tu and M. Tatar. Aging Cell 2, 327–333. https://doi.org/10.1046/j.1474-9728.2003.00064.x

65. Vijendravarma, R.K., Kawecki, T.J., 2015. Idiosyncratic evolution of maternal effects in response to juvenile malnutrition in Drosophila. J. Evol. Biol. 28, 876–884. https://doi.org/10.1111/jeb.12611

66. Vincent, C.M., Sharp, N.P., 2014. Sexual antagonism for resistance and tolerance to infection in Drosophila melanogaster. Proceedings of the Royal Society B: Biological Sciences 281, 20140987. https://doi.org/10.1098/rspb.2014.0987

67. Ye, Y.H., Chenoweth, S.F., McGraw, E.A., 2009. Effective but Costly, Evolved Mechanisms of Defense against a Virulent Opportunistic Pathogen in Drosophila melanogaster. PLOS Pathogens 5, e1000385. https://doi.org/10.1371/journal.ppat.1000385

68. Zuk, M., McKean, K.A., 1996. Sex differences in parasite infections: Patterns and processes. International Journal for Parasitology 26, 1009–1024. https://doi.org/10.1016/S0020-7519(96)80001-4

